# Frizzled1 and Frizzled2 are not redundant for competitive survival under low-Wingless levels in the developing *Drosophila* wing epithelium

**DOI:** 10.1101/2024.12.04.626737

**Authors:** Swapnil Hingole, Kritika Verma, Varun Chaudhary

## Abstract

In the *Drosophila* wing epithelium, canonical Wnt signaling is activated by the gradient of secreted Wingless protein (Wnt1 homolog), which interacts redundantly with the Frizzled1 and Frizzled2 receptors. While sharing overlapping functions, these receptors also have distinct non-canonical roles and exhibit differential expression patterns along the Wingless gradient. Moreover, Frizzled2, unlike Frizzled1, is thought to be essential for sustaining low-level Wingless signaling and promoting cell survival in the absence of the ligand. This raises the possibility of the two receptors acting differently along the Wingless gradient. In this study, we investigated the role of these receptors in cell survival across varying Wingless levels. We find that the loss of Frizzled2 in cells at a distance from the Wingless-producing cells—where Wingless levels are low— leads to competitive elimination of cells. In contrast, Frizzled1 is dispensable for cell survival, regardless of distance from the Wingless source. Our findings show that Frizzled2 is essential for competitive cell survival under low-Wingless conditions and the two receptors are not equally redundant across the Wingless concentration gradient, providing insight into a mechanism for spatial and temporal precision in developmental signaling.

## INTRODUCTION

In developing tissues, several cellular processes, including cell survival, are regulated by signaling molecules. Many of these molecules are released from localized sources and travel several cell distances to activate signaling. However, as cells divide and change positions relative to these sources, the ligand levels and consequently the survival signals available to them also change. Despite this challenge faced by the dividing cells, tissue development achieves remarkable robustness. This is largely due to the precise coordination between signal activation and feedback regulation (Kicheva and Briscoe, 2023).

A notable example of this robustness is the development of the *Drosophila* wing epithelium, which is regulated by the activity gradient of a secreted ligand called Wingless (Wg; Wnt1 homolog), besides several other signaling molecules. The Wg protein is released from a narrow strip of cells at the dorsal-ventral (DV) boundary (Couso et al., 1993; Strigini and Cohen, 2000; Williams et al., 1993), forming a symmetric gradient that activates signaling by interacting with the redundantly acting Frizzled1 (Fz1) and Frizzled2 (Fz2) receptors (Bhanot et al., 1999; Chen and Struhl, 1999; Kennerdell and Carthew, 1998; Müller et al., 1999). This signaling subsequently induces the expression of negative feedback regulators like Frizzled3 and Naked cuticle (Sato et al., 1999; Zeng et al., 2000), while concurrently repressing positive regulators such as Fz2 and the co-receptor Arrow (Cadigan et al., 1998). These feedback regulators assist in maintaining the appropriate levels of signaling.

Previously, it was shown that a steep difference in Wnt signaling activity between neighboring cells triggered a fitness-sensing mechanism known as cell competition, whereby cells with lower Wnt signaling levels are marked as “losers” and are eliminated by neighboring “winner” cells exhibiting high Wnt activity (Vincent et al., 2011). Interestingly, however, when a steep difference in Wg ligand levels between cells is generated by artificially tethering endogenous Wg to the membranes, thereby restricting its activity to juxtacrine mode, normal wing patterning is retained (Alexandre et al., 2014). We have previously shown that in the wing epithelium expressing only membrane-tethered Wg, higher levels of Fz2 in cells outside the range of tethered Wg maintain low-level expression of Wnt target genes (Chaudhary et al., 2019). Moreover, loss of Fz2—but not Fz1—resulted in the elimination of these “beyond-tethered-Wg” cells, suggesting a non-redundant role for Fz2 in cell survival in the absence of ligand. However, while *fz2* mutants in flies expressing tethered Wg exhibit severe lethality (Chaudhary et al., 2019), *fz2* mutants in otherwise flies with Wg show only a mild developmental delay, ultimately resulting in normally patterned wings (Chen and Struhl, 1999; Chen et al., 2004). Thus, it remained unclear whether this non-redundant function of Fz2 is also required for the development of normal wing discs with Wg gradient, possibly in cells distant from the source receiving low levels of Wg.

In this study, we have analyzed the role of Fz2 in cell survival along the Wg gradient. We generated clones of cells, randomly across the wing epithelium, either harboring *fz2* loss-of-function mutation or expressing *fz2-RNAi* and tested their survival over time. While Fz1 and Fz2 remained redundant for signaling and cell survival close to the source of Wg, we observed that cells at long range depended on *fz2* for survival. Moreover, we observed that loss of *fz2* under low-ligand conditions or following ligand removal resulted in the ‘loser’ cell fate, triggering cell competition and subsequent elimination by their neighboring fitter cells. In summary, our work shows that the known redundancy between Fz1 and Fz2 is not supported under low-ligand conditions, highlighting variation in their function along the Wg gradient.

## RESULTS AND DISCUSSION

### Fz2 deficient clones show impaired clonal propagation under low-Wg conditions

We set out to analyze the effect of Fz2 loss on cellular fitness in the presence of the endogenous Wg gradient in the developing wing epithelium. To this end, we generated clonal populations of cells homozygous for the *fz2* loss-of-function mutation (*fz2^−/−^)* through a mitotic recombination approach utilizing heat-shock-mediated Flippase(Flp) expression (see materials and methods). Clones were induced at the early larval stage (48 hrs after egg laying) and their growth was analyzed at 48 hrs and 72 hrs after clone induction (ACI). The generation of mitotic clones enabled a direct comparison of the growth of *fz2^−/−^* clones (GFP-negative) with *fz2^+/+^* twin spots (double-GFP positive with both functional *fz2* alleles) generated in parallel. Here, we observed that the growth of *fz2^−/−^*clones normalized to the nearby twin spots was significantly reduced as compared to the growth of the control clones normalized to their respective twin spots, at both 48 hrs and 72 hrs ACI (Fig1A-A’, B-B’, and C). In parallel, we also analyzed the clonal growth of cells expressing *fz2-RNAi.* Consistent with the mutant clones, we observed that the relative number of cells expressing *fz2-RNAi* is reduced over time (FigS1A-E), indicating that *fz2* is required for proper tissue growth during wing disc development.

**Figure 1:**
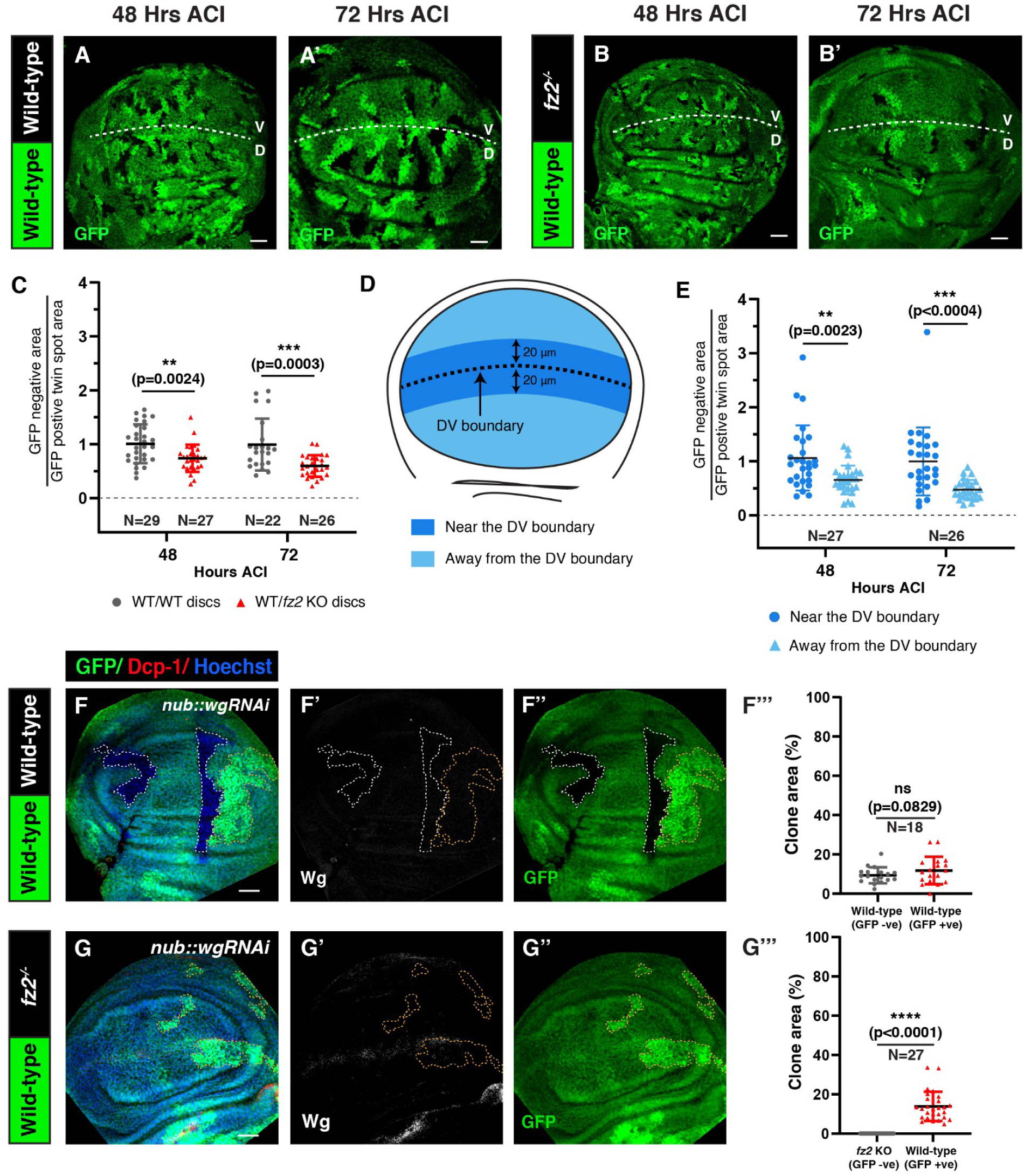
Fz2 deficient clones show impaired growth in low-Wg conditions. Images of wing discs showing growth of wild-type (A-A’) and *fz2* KO (B-B’) clones, 48, and 72 hours ACI. The white dotted line marks the DV boundary. (C) The graph represents the GFP-negative area normalized to the GFP-positive twin spot area per wing pouch region at 48, and 72 hrs ACI. (D) Schematic representation of wing imaginal disc showing the area with high Wg levels near the DV boundary (blue) and area with low Wg levels away from the DV boundary (light blue), (also see FigS2). (E) The graph represents the GFP-negative *fz2^−/−^* area normalized to the GFP-positive *fz2^+/+^*twin spot area near (high-Wg) and away (low-Wg) from the DV boundary at 48, and 72 hrs ACI. (F-F’’ and G-G’’) Images of wing discs expressing *wg-RNAi* in the wing pouch through *nub-Gal4*, harboring either wild-type *fz2^+/+^* clones (F-F’’) or *fz2^−/−^* clones (G-G’’) (72 hrs ACI). The white outline marks the GFP-negative area and the yellow outline marks the GFP-positive twin spot. Wg depletion is observed by Wg staining. Graphs in (F’’’, and G’’’) represent the percentage of area covered by GFP-positive twin spots compared to GFP-negative clones for respective genotypes. An Unpaired t-test (C), and a paired t-test (E, F’’’, and G’’’) were applied for statistical analyses. N values are mentioned in the graphs. Scale bar: 20 μm

Both cell proliferation and cell survival contribute to tissue growth and patterning. Previously, we have observed that in the wing epithelium expressing membrane-tethered Wg, Fz2 is essential for the survival of cells away from the reach of the ligand (Chaudhary et al., 2019), indicating that varying Wg levels could affect the redundancy between Fz1 and Fz2. Thus, we next asked if Fz2 could differentially affect cell survival along the Wg gradient. To test this, we established high-Wg and low-Wg regions based on the detectable range of endogenous Wg. Significant levels of Wg could be observed up to 20 μm distance on both sides of the DV boundary (FigS2A-B’’), which was defined as the high-Wg region, whereas the tissue beyond this point was considered as the low-Wg region (Fig1D and FigS2A-B’’). Careful analysis showed that the relative area of *fz2^−/−^* clones away from the DV boundary (low-Wg region) was reduced, compared with the *fz2^−/−^* clones near the DV boundary (high-Wg region) (Fig1B-B’, and E), indicating that Fz2 is essential under low-Wg conditions.

To further validate these findings, we analyzed the growth of *fz2^−/−^* clones in discs following the depletion of either the ligand Wg or the Wnt-trafficking protein Evenness interrupted (Evi; also known as Wntless), which is essential for the secretion of all lipid-modified Wnt proteins (FigS3A) in *Drosophila* (Bänziger et al., 2006; Bartscherer et al., 2006; Goodman et al., 2006). We have previously observed that in wing discs with reduced Evi levels, *fz2^−/−^* clones induced using *Ubx-Flp* showed lower signaling activity (Chaudhary et al., 2019), however, whether their survival was affected over time remained unknown. To this end, we generated *hs-Flp*-induced *fz2^−/−^* clones in wing discs expressing *wg-RNAi* or *evi-RNAi* in the entire pouch region via *nub-Gal4.* The control GFP-negative clones in either Wg or Evi depleted discs, observed at 72 hours ACI, showed no growth defects and were comparable to the twin spot (Fig1F-F’’’ and FigS3B-B’’’). In contrast, *fz2^−/−^* clones were not recovered at 72 hours ACI upon depletion of either Wg or Evi, whereas large twin spots could be observed (Fig1G-G’’’ and FigS3C-C’’’). Altogether, these results indicate that Fz2 is essential for the survival of cells under low-ligand conditions, observed at long-range from the DV boundary in normal wing discs. Furthermore, while Fz2 can also potentially interact with other *Drosophila* Wnts (Wu and Nusse, 2002), this function of Fz2 appears to be dependent on only Wg levels.

### Fz2 promotes cell survival under low-Wg conditions by providing a competitive advantage

Since Fz2 depletion caused increased cell death in membrane-tethered Wg discs (Chaudhary et al., 2019), we next tested the activation of cell death upon loss of Fz2 under low-Wg conditions. As expected, we observed higher levels of cell death, marked with anti-cleaved-death caspase-1 (Dcp-1) antibody in *fz2^−/−^* clones away from the DV boundary compared to the *fz2^−/−^* clones near the DV boundary (Fig2A-A’’’, B-B’’, and C-C’’). Whereas the control wild-type clones did not show an increase in cell death, regardless of the proximity to the DV boundary (FigS4A-A’’’, B-B’’, and C-C’’). Similarly, higher cell death was observed in *fz2-RNAi*-expressing clones compared with the control clones (Fig2D-D’’, E-E’’, and F). Moreover, cell death was higher in clones away from the DV boundary (Fig2E-E’’, and G), consistent with the clone growth data.

**Figure 2:**
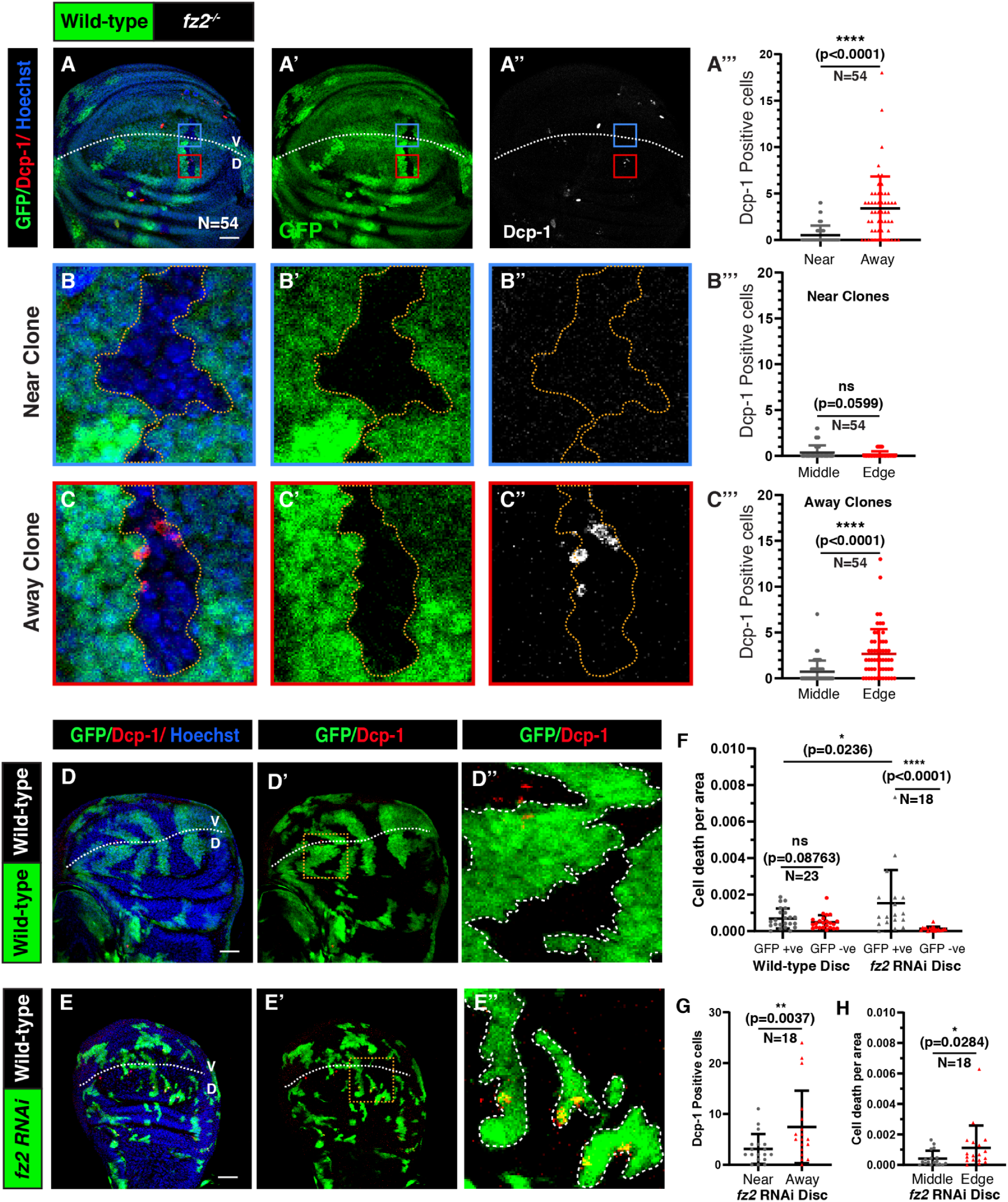
Fz2 deficient cells away from the DV boundary are eliminated via cell competition. (A-A’’) Cleaved Dcp-1 stained disc harboring GFP-negative *fz2*^−/−^ clones observed 72 hrs ACI. The graph in (A’’’) compares the number of cleaved Dcp-1 stained cells in the GFP-negative *fz2^−/−^*clones near (high-Wg) and away (low-Wg) from the DV boundary. (B-B’’) and (C-C’’) shows enlarged images of *fz2^−/−^* clones (marked by the yellow dotted line), near (blue box in A-A’’), and far from the DV boundary (red box in A-A’’), respectively. The graphs in (B’’’) and (C’’’) represent the cell death occurring in the middle of the *fz2^−/−^* clones compared to the edges of clones, near and away from the DV boundary, respectively. (D-E’’) Images of cleaved Dcp-1 stained disc harboring Actin Flipout Gal4 clones observed 72 hrs ACI, overexpressing *UAS-GFP* (D-D’’) and *UAS-fz2-RNAi* (E-E’’). The zoomed images show the area within the yellow box for respective images, and the white dotted line in the zoomed images marks the clone area. The graph in (F) represents Dcp-1 positive cells in the GFP-positive and GFP-negative area for control and *fz2-RNAi* discs. The graph in (G) represents the Dcp-1 positive cells in the *fz2-RNAi* clones near and away from the DV boundary. The graph in (H) represents the Dcp-1 positive cells per area in the middle and edges of the *fz2-RNAi* clones. A paired t-test (A’’’, B’’’, C’’’, G, and H) and an unpaired t-test (F) were applied for statistical analyses. N values are mentioned in the graphs. Scale bar: 20 μm

The wing epithelial cells grow in a competitive environment, where cells harboring defects that reduce cellular fitness are identified as “losers” by the neighboring fitter cells (Morata and Ripoll, 1975; Simpson, 1979; Simpson and Morata, 1981). This leads to the elimination of loser cells via apoptosis in a contact-dependent manner (Baker, 2020; de la Cova et al., 2004; Moreno and Basler, 2004; Moreno et al., 2002; van Neerven and Vermeulen, 2023). Since higher cell death is observed in both the *fz2^−/−^* and *fz2-RNAi* clones at long-range, we hypothesized that *fz2-RNAi* and *fz2^−/−^* cells gain the loser status and are outcompeted by surrounding wild-type cells. Consistent with this, the cell death in the *fz2^−/−^*clones away from the DV boundary was found to be significantly higher at the edges of the clones (Fig2C-C’’’ and FigS4D-D’’), suggesting that the death is induced by the neighboring wild-type cells in a contact-dependent manner, a hallmark of cell competition. Similarly, the cell death observed in *fz2-RNAi* clones is also higher at the edges of the clones (Fig2E-E’’, and H). However, we did not observe the same for *fz2^−/−^* clones at short-range (Fig2B-B’’’) or in the control wild-type clones (FigS4A-C’’’). These results suggest that the Fz2 deficient cells in the low-Wg region of the wing imaginal disc are perceived as less fit and subjected to competitive elimination from the tissue. The loser status of Fz2 deficient cells may be attributed to reduced canonical signaling, which aligns with previous studies demonstrating that cells with impaired Wnt signaling are eliminated via cell competition (Giraldez and Cohen, 2003; Johnston and Sanders, 2003; Vincent et al., 2011).

Next, we sought to test the effect of Fz2 loss with induced low-Wg conditions throughout the wing disc. To this end, we knocked down Fz2 in *evi^2^* flies (FigS5A). Initially, we depleted Fz2 in the posterior compartment of homozygous *evi^2^* (null mutant) discs, using *hh-Gal4.* However, continuous knockdown of Fz2 in *evi^2^* flies caused early larval lethality. Therefore, we temporally restricted the expression of *fz2-RNAi* in *evi^2^* flies for 48 hours, using temperature-sensitive Gal80. In these discs, we observed higher cell death in the posterior compartment as compared to the anterior control compartment (FigS5B-C’’’). Moreover, cell death was observed even closer to the DV boundary, contrary to the Fz2 knockdown in an otherwise wild-type background (FigS5B’’ and C’’). Collectively, these findings indicate that Fz2 is essential for cell survival under low- ligand conditions and cells lacking Fz2 in these conditions are designated as losers, leading to their elimination from the tissue via cell competition.

### Fz1 is dispensable for competitive survival along the Wg gradient

We next asked whether the impact of Fz2 depletion on cell survival could result from an overall decrease in Fz receptor levels rather than being a specific consequence of Fz2 loss. To address this, we aimed to determine whether a similar effect might also be observed following the depletion of Fz1 alone. We first generated the *fz1* KO clones and examined their growth. In contrast to the consequences of Fz2 loss, *fz1^−/−^*clones, observed at 72 hours ACI, showed similar growth as the twin spots (Fig3A-A’’’) and showed no effect on cell death regardless of their proximity to the DV boundary (Fig3A-A’’’, and B). Consistent with the findings of *fz1^−/−^* clones, the *fz1-RNAi* expressing clones also did not show any reduction in clone size compared to control clones (FigS6A-E) or a significant increase in cell death compared to control region (FigS6F-F’’’).

**Figure 3:**
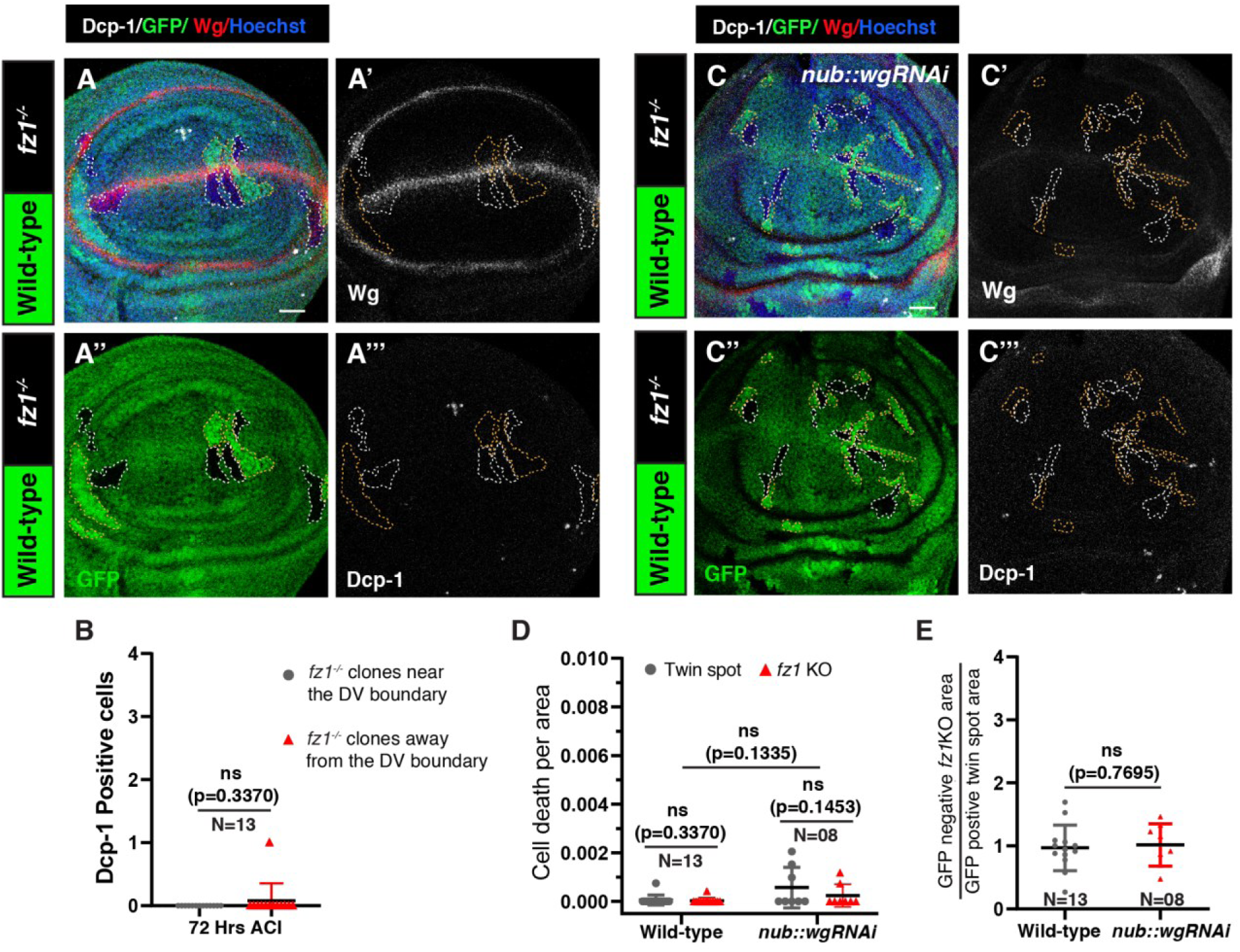
Cell survival is unaffected upon Fz1 loss under low-Wg conditions. (A-A’’’) Cleaved Dcp-1 and Wg stained disc harboring GFP-negative *fz1*^−/−^ clones (white outline) observed 72 hrs ACI. (B) The graph represents the number of Dcp-1 positive cells in *fz1*^−/−^ clones near (high-Wg) and away (low-Wg) from the DV boundary. (C-C’’’) Cleaved Dcp-1 and Wg stained disc harboring *fz1*^−/−^ GFP-negative clones (white outline) observed 72 hrs ACI in Wg knockdown disc. (D) The graph represents the cell death per area for *fz1*^−/−^ clones and wild-type *fz1^+/+^* twin spots in control and Wg knockdown discs. (E) Graph representing the ratio of *fz1* KO to wild-type twin spot clone areas in control and Wg knockdown discs. A Paired t-test (B, and D) and an unpaired t-test (D, and E) were applied for statistical analyses. N values are mentioned in the graphs. Scale bar: 20 μm

To further investigate the impact of Fz1 loss in a low-Wg context, we generated *fz1* KO clones in *wg-RNAi* discs. The *fz1^−/−^* clones survived in *wg-RNAi* discs and propagated comparably to the *fz1^−/−^* clones in otherwise wild-type discs, with no effect on cell death (Fig3A-A’’’, C-C’’’, D and E). Moreover, knocking down Fz1 in the posterior compartment of the *evi^2^*discs does not result in increased cell death (FigS7A-B’’’), unlike Fz2 knockdown (FigS5B-C’’’), suggesting that Fz1 is dispensable for cell survival under low-Wg conditions. Altogether, these results show that despite the known redundancy between Fz1 and Fz2 (Chen and Struhl, 1999), depleting Fz2, but not Fz1, could also reduce the survival of cells, albeit less severely than the concurrent loss of both Fz1 and Fz2 observed in previous studies (Giraldez and Cohen, 2003; Johnston and Sanders, 2003). Therefore, the functional redundancy between the two receptors is effective only under conditions of high ligand availability.

### Fz2 is required for maintaining proper wing size

So far, the clonal analysis shows that the redundancy between Fz1 and Fz2 varies along the concentration gradient of Wg. However, no significant phenotypes related to loss-of-Wnt signaling have been noted in the wing of *fz2* mutants (Chen and Struhl, 1999; Chen et al., 2004). It remains unclear whether this function of Fz2 is important for wing development. Thus, we performed a careful reassessment to determine whether *fz2* mutants have defects in the wing size.

To this end, we analyzed transheterozygous mutants carrying *fz2^C1^*and *Df(3L)fz2* (a deletion affecting *fz2* (Bhanot et al., 1999; Chen and Struhl, 1999)), to minimize potential background effects. We observed that the wings of transheterozygous *fz2^C1^*/*Df(3L)fz2* mutants are ~9-10% smaller compared to the control wild-type or heterozygous *fz2^C1^*/+ or *Df(3L)fz2/+* flies (Fig4A-A’). Additionally, these transheterozygous *fz2^C1^*/*Df(3L)fz2* mutants showed a significant delay in development (Fig4B), consistent with previous reports (Chen and Struhl, 1999). Although we cannot rule out the possibility that this developmental delay is due to the loss of Fz2 in other tissue(s), it is noteworthy that the wings failed to achieve their proper size despite this delay.

**Figure 4:**
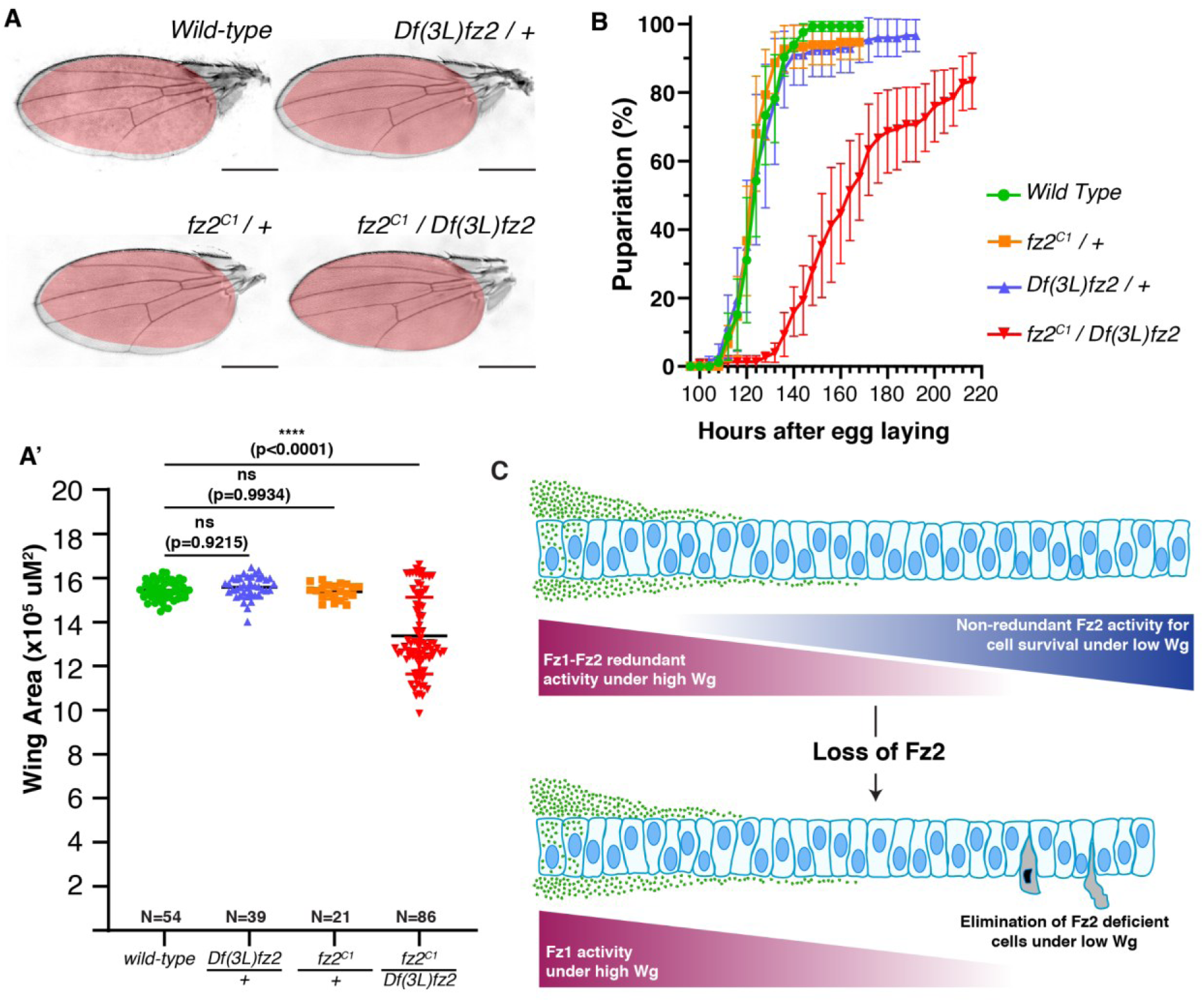
*fz2* mutants show reduced wing size and delayed development. (A-A’) Wing size of wild-type flies, heterozygous *fz2^C1^/+* and *Df(3L)fz/+* flies, and transheterozygous *fz2* knockout flies. (A’) The graph represents the average female wing size of the wild-type, *Df(3L)fz/+, fz2^C1^/+,* and *fz2* transheterozygous flies. A one-way ANOVA with Dunnett’s test was applied for statistical analysis. N values are mentioned in the graphs. (B) The graph shows pupariation timings of the wild-type, *fz2* heterozygous (*fz2^C1^/+*, and *Df(3L)fz/+*), and *fz2* transheterozygous animals. 150 larvae across five experiments for each genotype were used. (C) Wing development is compromised in the absence of Fz2. The cells in the high Wg region are unresponsive to Fz2 loss due to redundancy with Fz1. The cells in the low Wg region depend on high Fz2 levels for survival and are subjected to cell competition-mediated elimination upon losing Fz2. Scale bar: 500 μm

Overall, our results demonstrate that cells distal to the Wg source, which are exposed to low-Wg levels, depend on the positive feedback regulator Fz2 for their survival (Fig4C). The lack of redundancy at low-ligand levels is consistent with the higher affinity of Fz2 for Wg compared to Fz1 (Boutros et al., 2000; Rulifson et al., 2000; Wu and Nusse, 2002). Past studies have shown that overexpression of Fz2 leads to the stabilization of Wg over the receiving cells, protecting it from degradation (Baeg et al., 2004; Cadigan et al., 1998; Eldar et al., 2003). Thus, raising a possibility that reduced survival of Fz2 depleted cells at long-range may result from reduced Wg levels due to increased degradation. However, contrary to overexpression studies, extracellular Wg levels were shown to be unaffected in *fz2* mutant clones (Han et al., 2005), while the *fz1-fz2* double mutant clones show a mild increase in extracellular Wg levels (Baeg et al., 2004; Han et al., 2005; Piddini et al., 2005). Thus, the activity of Fz2 in regulating cell survival under low-ligand conditions may be governed by other, yet unidentified mechanisms. Furthermore, it remains to be determined whether the co-receptor Arrow, which is required for canonical activity and is a positive feedback regulator (Cadigan et al., 1998), could support Fz2 in this function.

Our data suggest that Fz2 levels affect competitive survival, potentially buffering the fluctuating levels of Wg activity observed in the Fz2-deficient cells by eliminating them from the tissue, leading to proper maintenance of signaling in the rapidly growing wing epithelia. Consistent with this, past studies in Zebrafish embryos have shown that the variability in Wnt gradient activity is corrected by cell competition-mediated elimination of cells with aberrant signaling, maintaining developmental robustness (Akieda et al., 2019). These findings also suggest that the redundant functions of Fz1 and Fz2 depend not only on the intrinsic properties of the receptors but also on buffering mechanisms like cell competition. Given that, similar to *Drosophila*, various organisms—including mice and humans—possess multiple genes encoding different Frizzled receptors with known redundancies within their subfamily (Wang et al., 2016), it would be intriguing to explore whether analogous cellular buffering mechanisms are triggered to compensate for the loss of these receptors.

## MATERIALS AND METHODS

### Drosophila genetics

The following stocks were used: *Actin5C-FRT-CD2-FRT-Gal4* (BDSC, 4779), *hs-Flp* on 1st chr., *UAS-GFP* on 3rd chr. (gifts from A. Teleman, German Cancer Research Center, Heidelberg), *UAS-fz2-RNAi* (KK-ID 108998), *hh-Gal4* 3rd chr. (Tanimoto et al., 2000), *tub-Gal80ts* (BL7108), *UAS-fz1-RNAi* (KK-ID 105493), *fz2^C1^ ri FRT2A (Chen and Struhl, 1999)*, *Ubi-GFP FRT2A* (BL1626), *fz^P21^ FRT80B* (Jones et al., 1996), *Ubi-GFP FRT80B* (BL1620), *FRT2A* (BDSC, 1997), *Df(3L)fz2* (BDSC, 6754), *evi^2^*(German Cancer Research Center, Heidelberg (Bartscherer et al., 2006)), *UAS-evi-RNAi* (KK-ID 103812), *UAS-wg-RNAi* (KK-ID 104579), *nub-Gal4* (Calleja et al., 2000). Detailed genotypes are mentioned in the supplemental information.

All crosses were reared on standard culture medium at 25°C, except where specifically mentioned.

### Antibodies

Larval wing imaginal discs were stained using the following antibodies: Rabbit anti-cleaved Dcp-1 (1:300, Cell Signaling Technology), rat anti-Fz2 (1:300, (Chaudhary et al., 2019)), mouse anti-Wg (1:50, Developmental Studies Hybridoma Bank (DSHB)), rat anti-Ci (1:50, DSHB). Secondary antibodies used for fluorescent labeling were Alexa-405, Alexa-488, Alexa-568, Alexa-594, Alexa-647 (Invitrogen) at 1:500 dilutions and Hoechst 33342, H3570 (1:1000, Invitrogen).

### Immunostaining

Larvae of desired genotypes were dissected in Phosphate-buffered saline (PBS), and head complexes with wing imaginal discs were separated, followed by fixation with 4% paraformaldehyde (PFA) for 30 minutes at room temperature. Subsequently, samples were permeabilized with PBS-T (0.2% Triton in 1XPBS), followed by blocking using BBT (0.1% BSA in PBS-T) and overnight incubation in the primary antibody at 4°C. The following day, the primary antibody was removed and samples were washed with PBS-T followed by incubation with fluorophore-conjugated secondary antibody for 90 minutes at room temperature. After removing the secondary antibody, a few PBS-T washes were given to remove excess nonspecific staining. The wing discs were mounted in Vectashield (Vector Labs) mounting media. Staining and microscopy conditions for the samples used were identical. Wing discs are oriented with the ventral up and anterior left.

### Image acquisition and processing

Images of fixed samples were acquired using the 40x oil objective on the Olympus (FV3000) confocal microscope, and Olympus Spinning Disc microscope, with each slice (z-stack) equivalent to 1μm. Images were processed using ImageJ (Fiji) and Adobe Photoshop CS6 v13.0. Figures were made in Adobe Illustrator (Adobe Illustrator CS6 Tryout version 16.0.0). Schematics and models were made in Adobe Illustrator (Adobe Illustrator CS6 Tryout version 16.0.0).

### Generation of clones

For generating flip-out and mitotic clones, the FLP (Flippase)/FRT (Flippase recognition target) system was used under a heat shock-driven promoter. Mitotic clones were induced by giving a heat shock of 37°C for 60 minutes at 48 hrs AEL (after egg laying), and larvae were then shifted to 25°C. Actin Flip-out Gal4 (AFG) clones were induced by giving a heat shock of 37°C for 15 minutes at 48 hrs AEL (after egg laying), and larvae were then shifted to 25°C. Larvae were dissected at 48, and 72 hrs ACI.

### Pupariation assay

Early larvae (~48 hrs after egg laying) of the desired genotype were selected using tubby or GFP balancer chromosomes. Thirty larvae of each genotype were transferred into fresh food vials and kept at 25°C under normal laboratory conditions. The number of larvae pupariated for each genotype was recorded at 4 hr intervals. A minimum of 150 larvae were used across five independent experiments performed for each genotype.

### Quantification and analysis

For all the analyses of wing discs, only the wing pouch area was considered. The area of clones and pouch region was measured using ImageJ. The area of mitotic clones was analyzed by measuring the total clone area and normalizing it with the total twin spot area for each sample (Fig1C, E, and Fig3E). Separately, the area for mitotic clones, and twin spots in Wg and Evi knockdown discs (Fig1F’’’, G’’’, FigS3B’’’, and C’’’) and Actin Flip-out Gal4 clones (FigS1E, and FigS6E) were analyzed by calculating the percentage of area covered by clones normalized to the wing pouch area. The cell death was analyzed by measuring the cleaved Dcp-1 positive cells in the area of interest (Fig2A’’’, B’’’, C’’’, F, G, and H, Fig3B, and D, FigS3A’’’, B’’’ and C’’’, and FigS6F’’’).

Wing areas of female flies were measured for individual samples for respective genotypes (Fig4A’). Pupated larvae for each time across five replicates were recorded and averaged to determine pupariation timing for respective genotypes (Fig4B). MS Excel was used to record all the data and perform the necessary analyses. GraphPad Prism 8 was used to make the graphs and perform statistical analyses.

## ACKNOWLEDGMENTS

We thank M. Boutros, G. Struhl, A. Teleman, the Bloomington Stock Center, and the Vienna *Drosophila* Research Center (VDRC) for *Drosophila* strains and reagents. We also thank D. Strutt for the Fz2 antibody. We are grateful to IISER Bhopal for the fly facility and the DST-FIST facility for the confocal microscopy. We thank V.C. lab members for the discussions on the project and manuscript.

## AUTHOR CONTRIBUTIONS

Conceptualization: All authors; Investigation: SH and KV; Formal analysis and methodology: SH and KV; Validation: SH and KV; Writing - original draft preparation: SH and VC; Writing - review and editing: All authors; Supervision: VC; project administration: VC; funding acquisition: VC. All authors have read and agreed to the published version of the manuscript.

## FUNDING INFORMATION

This work was funded by a Department of Biotechnology (DBT) grant to VC (BT/PR34467/BRB/10/1831/2019). VC’s laboratory is also supported by intramural funds from IISER Bhopal. SH and KV were supported by senior research fellowships from DBT.

## CONFLICT OF INTEREST STATEMENT

The authors declare no conflicts of interest.

## Notes

### Competing Interest Statement

The authors have declared no competing interest.

