## Supplemental Data for "Frizzled1 and Frizzled2 are not redundant for competitive survival under low-Wingless levels in the developing *Drosophila* wing epithelium"

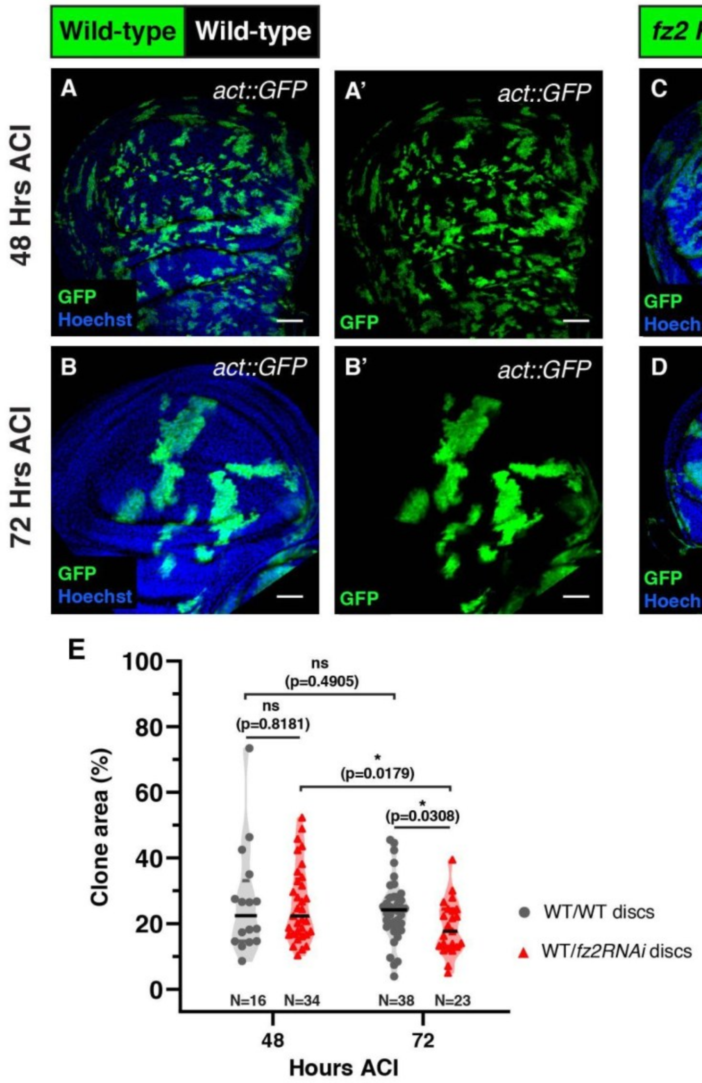


**Figure S1:** **Fz2 knockdown clones are gradually eliminated from the tissue.**

Representative images of wing discs showing the *hs-FLP* induced Actin Flipout Gal4 clones overexpressing control *UAS-GFP* (A-B') and *UAS-fz2-RNAi* (C-D') at 48, and 72 hrs ACI. (E) The graph represents the progression of the total clone area over time in the wing pouch region (%) for control *UAS-GFP* and *UAS-fz2-RNAi* clones. An unpaired t-test was used for statistical analysis, N values are mentioned in graph E. Scale bar: 20 µm

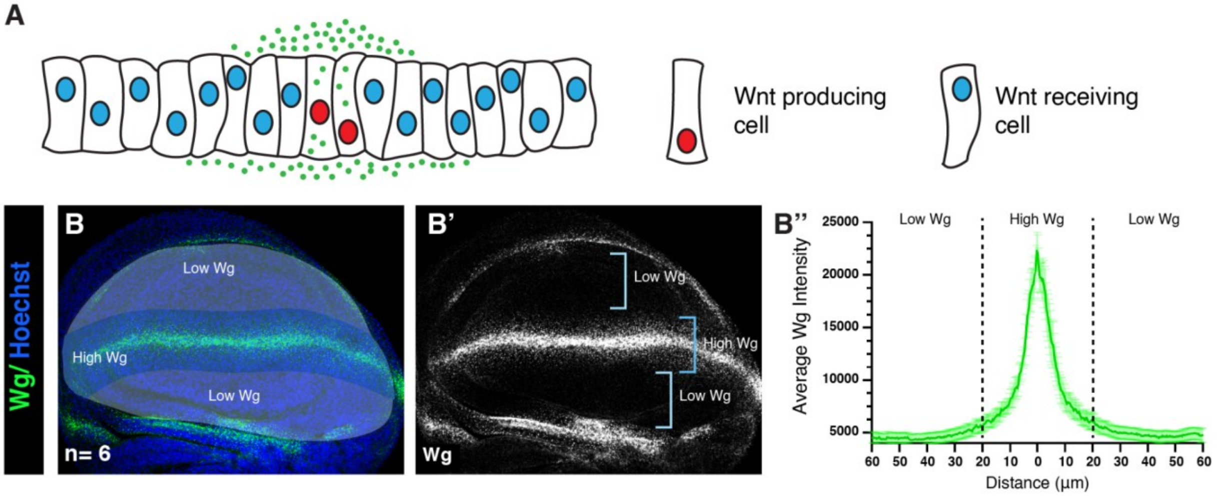


**Figure S2: Wg gradient in the wing disc.**

(A) Schematic representation of the orthogonal view of wing imaginal discs with Wg-producing cells (marked with the red nucleus) along the dorsoventral boundary from where the Wg is secreted to form a concentration gradient to reach the receiving cells (marked with the blue nucleus). (B-B') Wg stained wing imaginal discs with high levels (marked with blue bracket) and low levels (marked with light blue bracket) of Wg along the DV boundary. (B'') The graph represents the average levels of Wg across the multiple wing pouches with the DV boundary at the middle.

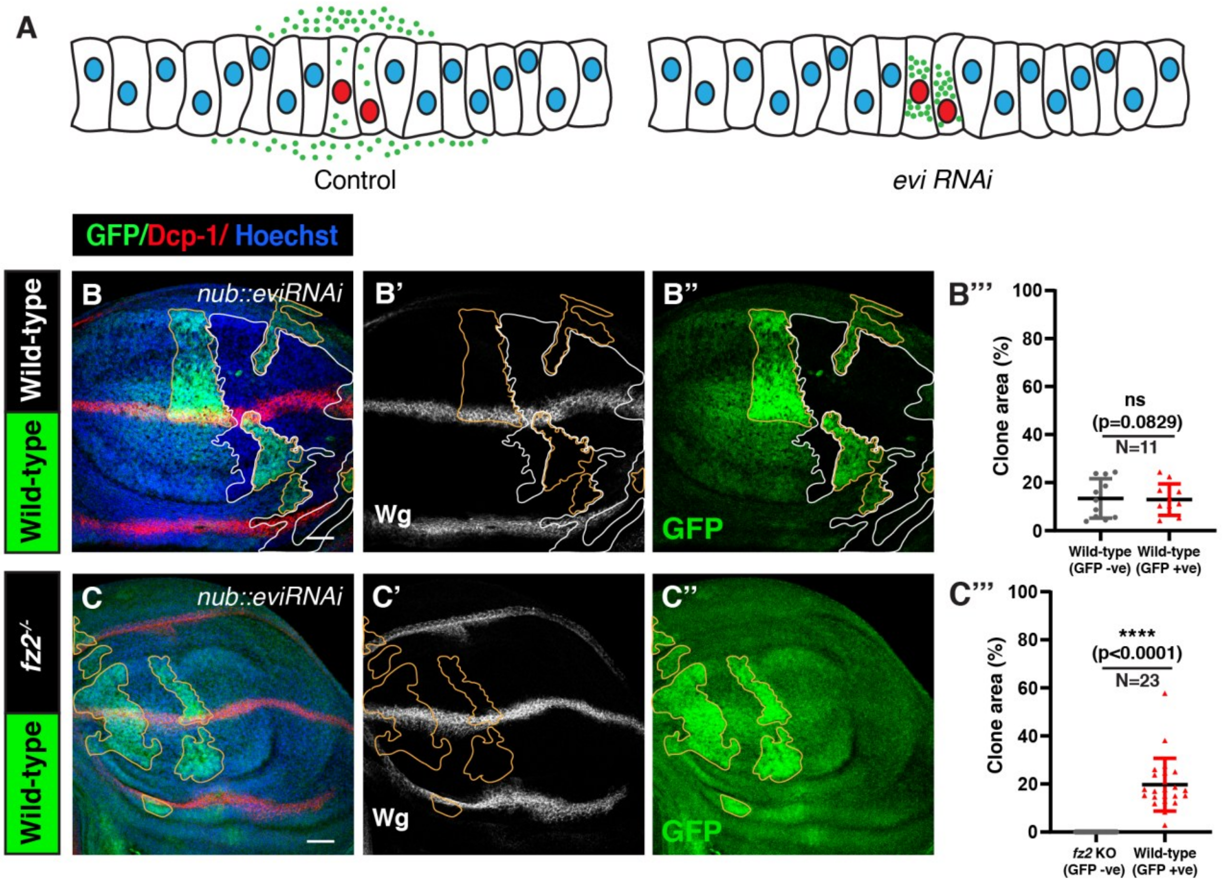


**Figure S3: *fz2* mutant clones are eliminated in the absence of secreted Wnts.**

(A) Schematic representation of Wg distribution in the presence of Evi (control disc) and the absence of the Evi (*evi-RNAi* disc). (B-B'' and C-C'') Images of wing discs expressing *evi-RNAi* in the wing pouch through *nub-Gal4*, harboring either wild-type $fz2^{+/+}$ clones (B-B'') or $fz2^{-/-}$ clones (C-C'') (72 hrs ACI). The white outline marks the GFP-negative area and the yellow outline marks the GFP-positive twin spot. Evi depletion is observed by the accumulation of Wg. Graphs in (B''', and C''') represent the percentage of area covered by GFP-positive twin spots compared to GFP-negative clones for respective genotypes. A paired t-test was applied for statistical analysis. N values are mentioned in the graphs. Scale bar: 20 μm

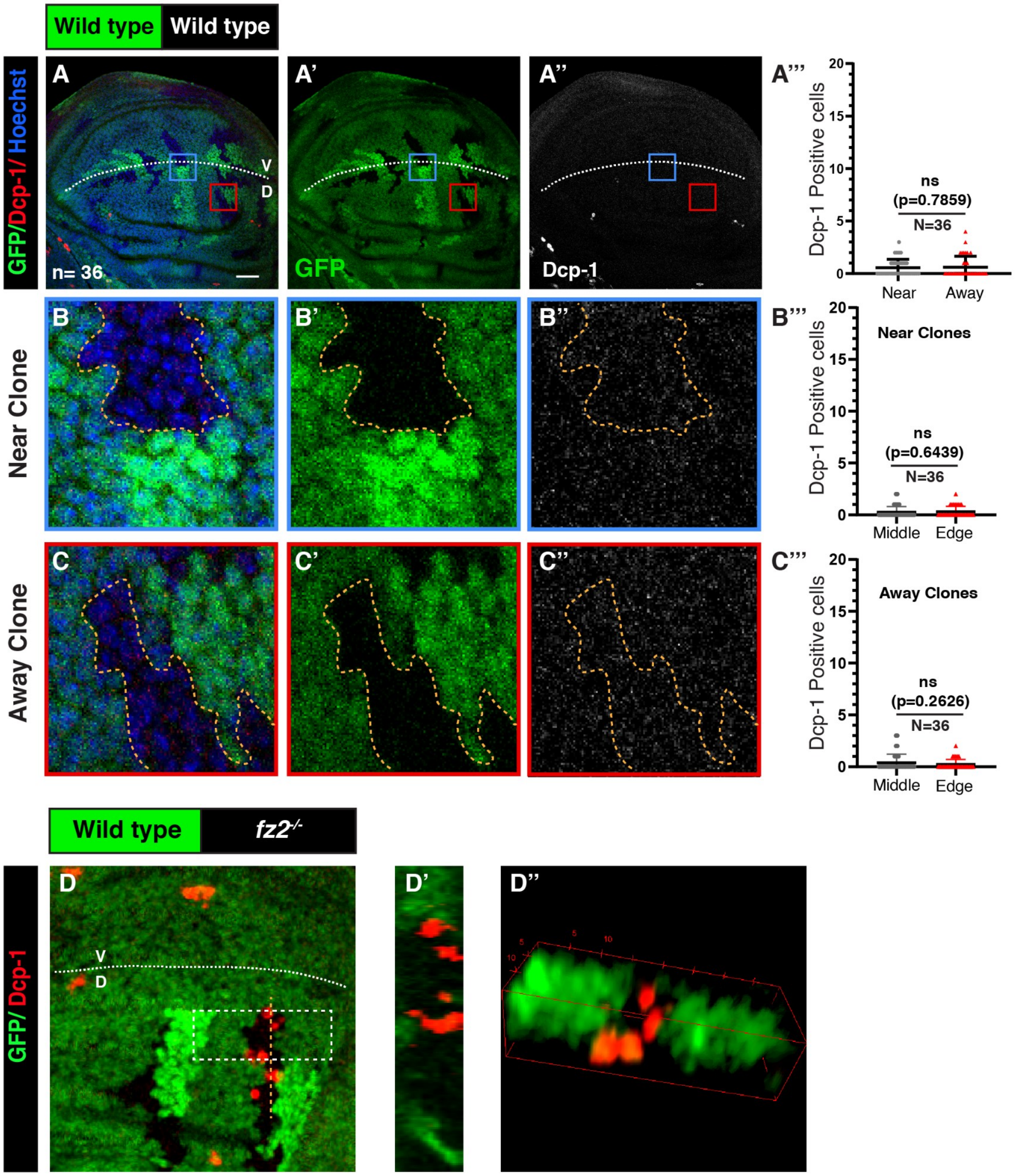


**Figure S4: Cell death is higher at the edges of *fz2* mutant clones present far from the DV boundary.**

**Figure S4: Cell death is higher at the edges of *fz2* mutant clones present far from the DV boundary.**

(A-A'') Cleaved Dcp-1 stained control disc harboring wild-type GFP-negative *fz2*$^{+/+}$ clones and wild-type GFP-positive *fz2*$^{+/+}$ twin spots observed 72 hrs ACI. (A''') The graphs represent the number of cleaved Dcp-1 stained cells in the wild-type GFP-negative *fz2*$^{+/+}$ clones near (high-Wg) and away (low-Wg) from the DV boundary. (B-B'', and C-C'') show enlarged images of GFP-negative *fz2*$^{+/+}$ clones (marked by the yellow dotted line), near (blue box in A-A''), and away from the DV boundary (red box in A-A''). The graphs in (B''') and (C''') represent the cell death occurring in the middle of the wild-type *fz2*$^{+/+}$ GFP-negative clones compared to the edges of the wild-type *fz2*$^{+/+}$ GFP-negative clones near and away from the DV boundary, respectively. (D) Cleaved Dcp-1 stained disc harboring *fz2* KO clones observed at 72 hrs ACI. (D') The transverse section along the yellow dotted line in (D). (D'') Volume representation of *fz2* KO clone in the white box in F. A paired t-test was applied for statistical analysis. Scale bar: 20 µm

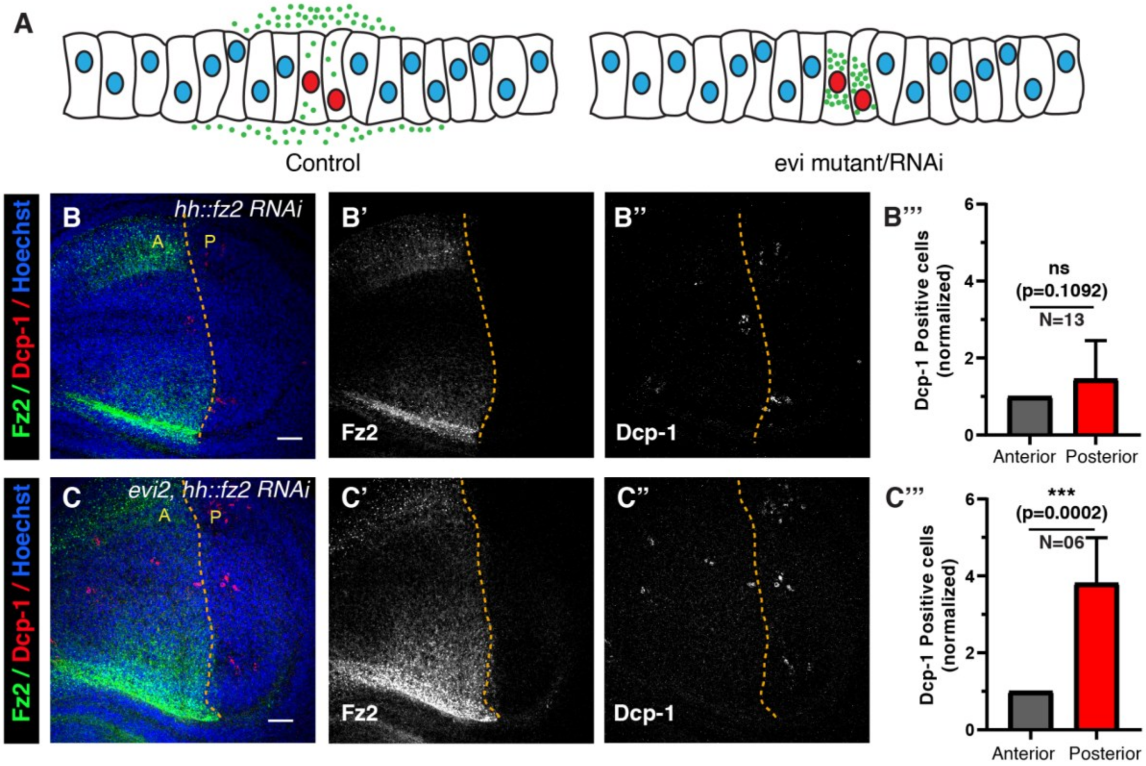


**Figure S5: Fz2 is necessary for cell survival in the absence of secreted Wnt/Wg.**

(A) Schematic representation of Wg distribution in the presence of Evi (control disc) and the absence of the Evi (*evi* mutant or RNAi disc). Control (B-B'') and *evi*$^{2}$ discs (C-C'') expressing *fz2*-RNAi in the posterior compartment for 48 hrs via *tubGal80*$^{ts}$*; hhGal4* and stained for Fz2 and Dcp-1. The yellow dotted line separates anterior and posterior compartments, marked by the Fz2 staining. (B''' and C''') The graphs represent normalized Dcp-1-positive cells in the anterior and posterior compartments for the respective genotypes. A paired t-test was applied for statistical analysis. N values are mentioned in the graphs. Scale bar: 20 μm

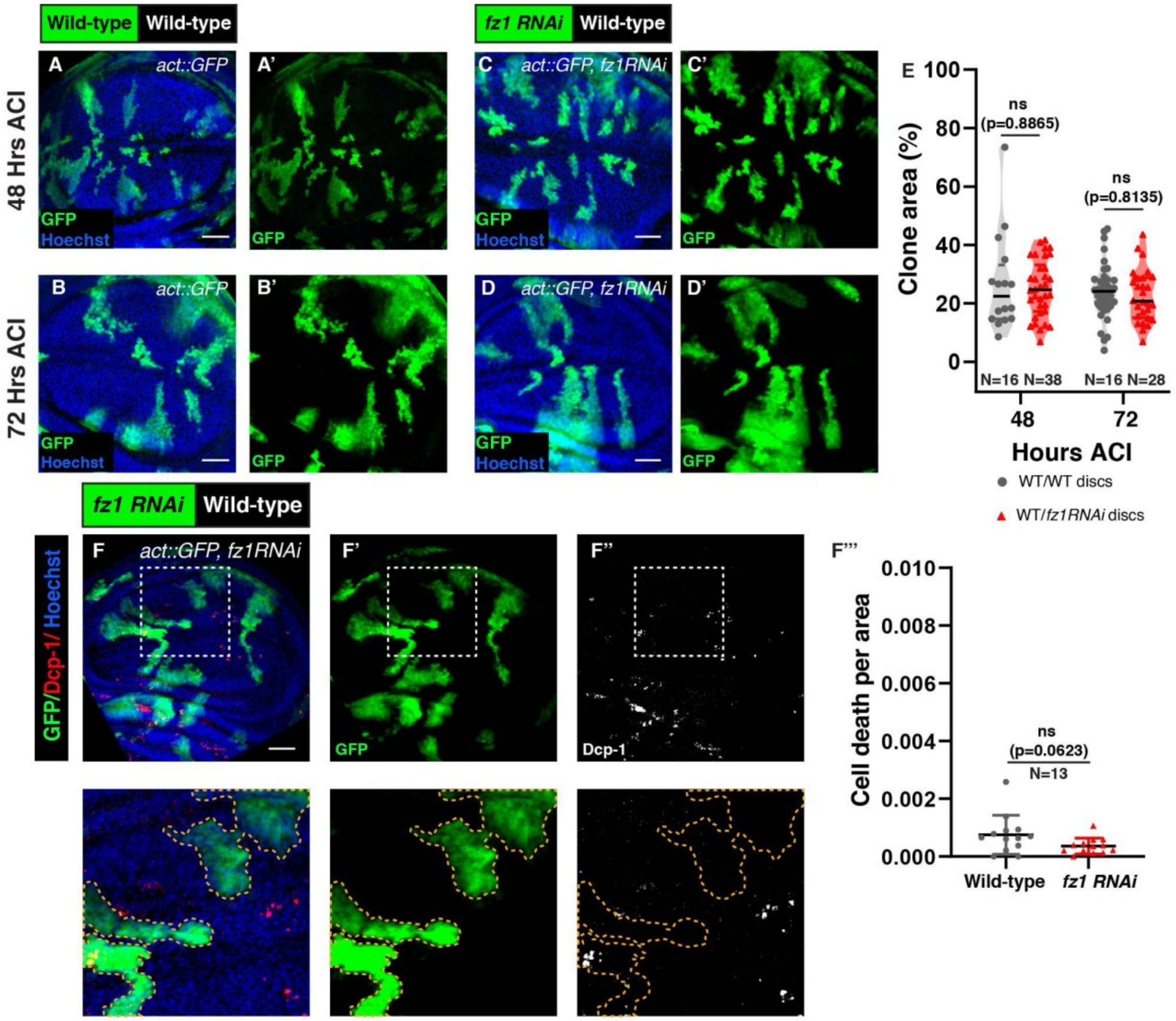


**Figure S6: Loss of Fz1 does not affect cell survival.**

Representative images of wing discs showing growth of control Actin Flip-out Gal4, *UAS-GFP* clones (A-B'), and *UAS-fz1-RNAi* clones (C-D') at 48 and 72 hours ACI. (E) The graph represents the progression of the total clone area over time in the wing pouch region (%) for control *UAS-GFP,* and *UAS-fz1-RNAi* clones. The zoomed images show the area within the white dotted box for respective images, and the yellow dotted line in the zoomed images marks the clone area. The graph in (F''') represents the Dcp-1 positive cells per area in control and *fz1-RNAi* clones. An unpaired t-test was used for statistical analysis, N values are mentioned in graphs. Scale bar: 20 μm

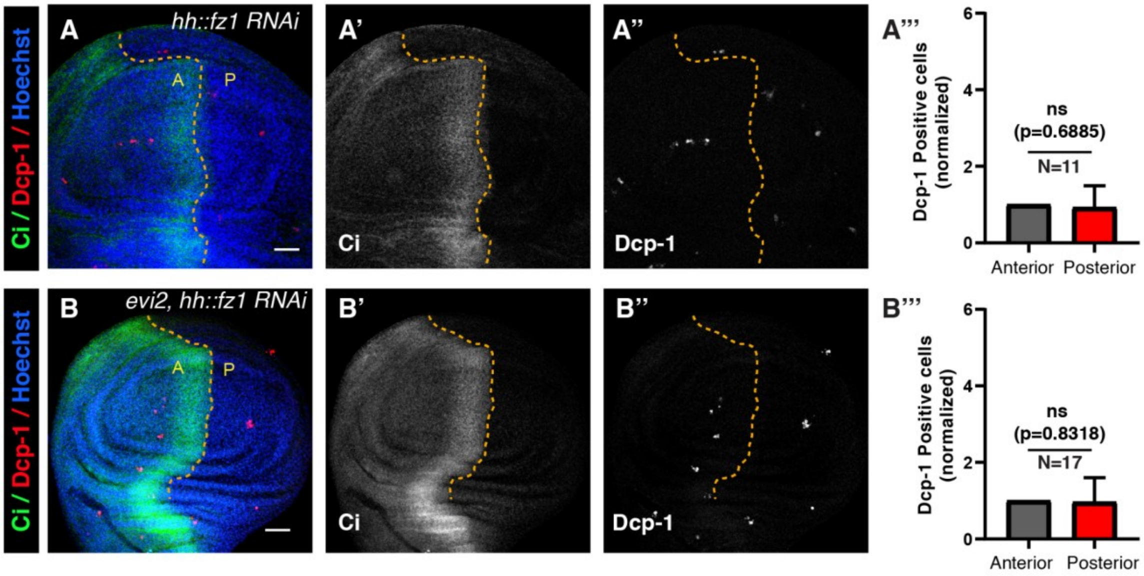


**Figure S7: Fz1 knockdown does not increase cell death in *evi*$^2$ discs.**

Control (A-A'') and *evi*$^2$ discs (B-B'') expressing *fz1*-RNAi in the posterior compartment for 48 hrs via *tubGal80*$^{ts}$*; hhGal4* and stained for Ci and cleaved Dcp-1. The yellow dotted line separates anterior and posterior compartments, marked by the Ci staining. (A'''-B''') The graphs represent normalized Dcp-1-positive cells in the anterior and posterior compartments for the respective genotypes. The paired t-test was applied for statistical analysis. N values are mentioned in the graphs. Scale bar: 20 μm

**SUPPLEMENTAL MATERIALS AND METHODS**

***Drosophila* genotypes**

The following genotypes were used in this study.

1A- 1A’: *hs-FLP / + ; + / + ; FRT2A / UbiGFP FRT2A*

1B- 1B’: *hs-FLP / + ; + / + ;* $fz2^{C1}$ *ri FRT2A / UbiGFP FRT2A*

1F-1F’’: *hs-FLP / + ; nubGal4 / UAS-wg-RNAi (KK) ; FRT2A / UbiGFP FRT2A*

1G-1G’’: *hs-FLP / + ; nubGal4 / UAS-wg-RNAi (KK) ;* $fz2^{C1}$ *ri FRT2A / UbiGFP FRT2A*

2A-2A’’: *hs-FLP / + ; + / + ;* $fz2^{C1}$ *ri FRT2A / UbiGFP FRT2A*

2D-2D’’: *AFG / hs-FLP ; +/+ ; UAS-GFP / +*

2E-2E’’: *AFG / hs-FLP ; UAS-fz2-RNAi (KK) / + ; UAS-GFP / +*

3A-3A’’’: *hs-FLP / + ; + / + ;* $fz1^{P21}$*FRT80B / UbiGFP FRT80B*

3C-C’’’: *hs-FLP / + ; nubGal4 / UAS-wg-RNAi (KK) ;* $fz1^{P21}$*FRT80B / UbiGFP FRT80B*

4A: *w1118* (upper left) *, Df(3L)fz2 / +* (upper right) *,* $fz2^{C1}$ *ri FRT2A / +* (lower left) *,* $fz2^{C1}$ *ri FRT2A / Df(3L)fz2* (lower right)

S1A-S1B’: *AFG / hs-FLP ; +/+ ; UAS-GFP / +*

S1C-S1D’: *AFG / hs-FLP ; UAS-fz2-RNAi (KK) / + ; UAS-GFP / +*

S2B-S2B’: *w1118*

S3B-S3B’’: *hs-FLP / + ; nubGal4 / UAS-evi-RNAi (KK) ; FRT2A / UbiGFP FRT2A*

S3C-S3C’’: *hs-FLP / + ; nubGal4 / UAS-evi-RNAi (KK) ;* $fz2^{C1}$ *ri FRT2A / UbiGFP FRT2A*

S4A-S4A’’: *hs-FLP / + ; + / + ; FRT2A / UbiGFP FRT2A*

S4D-S4D’’: *hs-FLP / + ; + / + ;* $fz2^{C1}$ *ri FRT2A / UbiGFP FRT2A*

S5B-S5B’’: *+ / + ; UAS-fz2-RNAi(KK) / tubGal80(ts) ; hhGal4 / +*

S5C-S5C’’: *+ / + ; UAS-fz2-RNAi(KK) / tubGal80(ts) ; hhGal4,* $evi^2$ */* $evi^2$

S6A-S6B’: *AFG / hs-FLP ; +/+ ; UAS-GFP / +*

S6C-S6D’: *AFG / hs-FLP ; UAS-fz1-RNAi (KK) / + ; UAS-GFP / +*

S6F-S6F’’: *AFG / hs-FLP ; UAS-fz1-RNAi (KK) / + ; UAS-GFP / +*

S7A-S7A’’: *+ / + ; UAS-fz1-RNAi(KK) / tubGal80(ts) ; hhGal4 / +*

S7B-S7B’’: *+ / + ; UAS-fz1-RNAi(KK) / tubGal80(ts) ; hhGal4,* $evi^{2}$ / $evi^{2}$
